# Challenges in predicting PROTAC-mediated Protein-Protein Interfaces with AlphaFold reveal a General Limitation on Small Interfaces

**DOI:** 10.1101/2024.03.19.585735

**Authors:** Gilberto P. Pereira, Corentin Gouzien, Paulo C. T. Souza, Juliette Martin

**Author notes:** **Corresponding Author,** Juliette Martin – Laboratoire de Biologie et Modélisation de la Cellule, CNRS, UMR 5239, Inserm, U1293, Université Claude Bernard Lyon 1, Ecole Normale Superieure de Lyon, 46 Allée d’Italie, 69364, Lyon, France. First authorship is shared between these authors (Gilberto P. Pereira, Corentin Gouzien).

## Abstract

Proteolysis Targeting Chimeras (PROTACs) are heterobifunctional molecules composed by ligands binding to a target protein and a E3-ligase complex, connected by a linker, that induce proximity-based target protein degradation. PROTACs are promising alternatives to conventional drugs against cancer. Predicting PROTAC-mediated complexes is often the first step for *in silico* PROTAC design pipelines. AlphaFold2 (AF2) revolutionized structural biology, enabling the prediction of multimeric protein structures. However, we previously noted that AF2 fails to predict PROTAC-mediated complexes.

Here, we investigate the potential causes of this limitation. We consider a set of 326 protein heterodimers orthogonal to the AF2 training set, and evaluate AF2 models focusing on the interface size and presence of interface ligand. Our results show that AF2-multimer predictions are sensitive to the size of the interface to predict even in the absence of ligands, with the majority of models being incorrect for the smallest interfaces. We also benchmark both AF2 and AF3 on a set of 28 PROTAC-mediated dimers and show that AF3 does not significantly improve upon the accuracy of AF2. The low accuracy of AF2 on complexes with small interfaces has strong implications for computational pipelines for PROTAC design, as these stabilize typically small interfaces, and more generally on any prediction task that involves small interfaces.

## Introduction

In the quest to develop innovative therapies, pharmaceutical companies and research laboratories invest billions of dollars every year into the different stages of drug design pipelines^1^. Recently, many of them have invested into the development of Proteolysis TArgetting Chimeras (PROTACS). PROTACs are heterobifunctional ligands composed of two small molecules connected by a linker region. One of the small molecules binds to a protein target and the other binds to a protein called E3 ligase, which is attached to the ubiquitination machinery^2,3^. This machinery is responsible for ubiquitination of proteins, which tags them for proteasomal degradation. The formation of a complex between the E3 ligase and the protein target *via* the PROTAC thus induces the degradation of the target. PROTAC-based approaches have been explored for a variety of therapeutic targets, including proteins intimately connected to cancer ^4–6^. These molecules have several advantages over small molecule-based drugs due to their catalytic mode of action. Due to it, the concentration at which a therapeutic effect is achieved tends to be lower than standard small molecule inhibitors, leading to a lower risk of side-effects ^7^. However, up until a few years ago, these molecules were designed serendipitously due to both a lack of structural information and limited successful applications of *in silico* tools.

Thanks to the effort undertaken the last few years, there now exists a wealth of structural, biochemical and activity data regarding several different PROTAC families which can be exploited toward the construction of computational pipelines for PROTAC design. The first step in these pipelines typically pertains to the prediction of the E3-ligase/target complexes, which can be achieved by a myriad of methods. While some groups have developed tools for this purpose^8–12^, they are typically not general because they require that the PROTAC molecule be known a priori. Within our group we developed a pipeline called PROTACability^13^, which bypasses this constraint and achieves satisfactory accuracy with minimal prior information. PROTACability was based on predictions of the E3 ligase/target interfaces produced by LightDock, a classical docking software^14^ .

The field of protein structure prediction has been deeply impacted by the AlphaFold method^15^, as well as protein-protein docking. Indeed, although not initially developed for protein-protein complex prediction, AlphaFold has been rapidly exploited by the community toward this goal, using hacks such as concatenating the sequence of several proteins using multi-glycine spacers ^16^ or modifying the input residue index ^17–19^ . AlphaFold-multimer was then specifically developed for protein assemblies^20^. These developments offer a new alternative for protein-protein docking. Recently, the new iteration of AlphaFold culminated into AlphaFold 3, the newest version which is able to take into consideration the information of ligands and model multimeric proteins accurately^20,21^. AlphaFold3 also differs from AlphaFold2 in several other aspects, including a smaller and simpler embedding step for the multiple sequence alignment (MSA), a new diffusion model in place of the AF2 structure module, early stopping during training and random noise sampling during inference time^22^. This results in an improved prediction of protein-protein complexes, particularly for antibody-antigen complexes, suggesting a lower dependency on co-evolutionary signal between entities.

During the development of PROTACability, we also considered using AlphaFold-multimer, with little success^13^. A recent benchmark of docking tools for PROTAC-mediated complexes confirm these poor quality results^13,23^. In this article, we explore further the reasons for this failure. We first use a test set of protein-protein heterodimers never seen by AlphaFold during the training phase to investigate the effect of the interface size and presence of ligand at the interfaces on the quality of the generated models. We then test the capacity of AlphaFold3 on the PROTAC-mediated interfaces on the relevant dimers, as well as the dimers in presence of the accessory proteins to better mimic the biological context of the interactions and reduce the search space. The benchmark on protein-protein heterodimers should help us to identify whether the difficulty with PROTAC-mediated complexes lies in (i) the interface size, (ii) the presence of a ligand, or (iii) the stabilisation of a non-natural complex resulting in the absence of any co-evolutionary signal. Our results point at an inherent limitation of AlphaFold-multimer on small interfaces in the general case, which compromises the prediction of PROTAC-mediated interfaces which are generally small. AlphaFold3 does not remedy this situation, even when the prediction context is provided. Nonetheless, as AF3 predictions were carried out without the presence of PROTACs in the available web server, it could be the case that AF3 full capabilities have not been explored.

## Material and Methods

### Dataset

A dataset of 28 PROTAC-mediated complexes was extracted from the PDB. These 28 cases represent the cases available at the time of the study and were chosen based on the following criteria: (i) they should contain at least the ternary complex (E3-ligase, protein target and PROTAC), (ii) resolution better than <4 Å, (iii) no missing residues at the protein-protein interface. In 26 cases, accessory proteins bound to the E3 ligase receptor are present. These accessory proteins are ignored in the prediction, unless otherwise stated.

A test set was extracted from the PDB as follows. All protein heterodimers (two different peptidic chains in the assembly) deposited after October 1st 2021 (training date of AlphaFold2 v3) were retrieved from the RCSB PDB website. The redundancy against the AlphaFold2 training set was removed using Foldseek^24^ (software version fc40ba1afdcb20602116907fdc73a47fd4c78615) against PDB sequences with a 30% sequence identity cutoff and an 80% sequence coverage cutoff (*easy-multimersearch --min-seq-id 0.3 --cov-mode 2 -c 0.8)*. Dimers with detectable sequence similarity to complexes acquired before 30 sept 2021 were removed. The intra-set redundancy was then reduced using cd-hit ^25,26^ (software version 4.8.1), with a 40% cutoff (step 1: redundancy reduction on a per-chain basis; step 2: addition of all chains of complexes selected in step 1, step 3: removal of remaining redundancy). Sequences shorter than 30 residues were removed. The resulting test set is composed of 340 heterodimers. Among these 340 dimers, 5 cases only were mediated by a ligand (see below).

Structures anterior to the AlphaFold training date were added to increase the number of ligand-mediated interfaces. We used the Dockground resource (https://dockground.compbio.ku.edu/bound/index.php)^27^ to build a set of non-redundant (<30% sequence identity) hetero-dimers X-ray structures with a resolution better than 3 Å deposited earlier than 30 sep 2021. A list of 20 ligand-mediated complexes was extracted from this set. The list of the complexes used throughout this study is given in **Table S1**.

### Interface size

The interface size was defined as the accessible surface area (ASA) of separated chains minus the ASA of the complex : *Δ = ASA(chain1) + ASA(chain2) − ASA(complex)*. ASA is computed using naccess ^28^ with default parameters (probe size=1.4 Å, Z slice = 0.05 Å).

Unless otherwise specified, hetero-atoms (i.e., non-standard residues, solvent and ligands) are excluded from the computation.

### Ligand-mediated interfaces

Ligand-mediated interfaces were identified by comparing the *ΔASA* computed with and without hetero-atoms, thus excluding peptidic ligands. Cases where the difference between *ΔASA* with and without hetero-atoms was greater than 10% were manually verified to exclude complexes with cross-links, modified residues at the interface, or non-specific ligands at the interface (such as sulfate ions, glycerol or PEG groups).

### AlphaFold 2

AlphaFold-Multimer calculations were carried out using ColabFold version 1.5.1^17,20,29^ using the alphafold2_multimer_v3 model with default parameters (i.e. paired+unpaired MSA, with model recycling). The best model was selected using the composite AF2 predicted quality score defined by 0.8 ipTM+0.2pTM^20^ . We will refer to this score as the AF2 confidence score. Among the 340 cases of the test set, AF2 was able to produce a model for 326 cases, including the 5 cases with ligand-mediated interface.

### AlphaFold 3

AlphaFold 3 predictions were run on the web server (https://alphafoldserver.com/) on the 6th of June 2024, solely for PROTAC cases. At that time, the webserver could only model common biological ligands, i.e., not PROTACs. Two experiments were carried out: in the first one, AF3 was provided with both sequences of the ligase and target (like AF2); in the second experiment we added the sequence of the accessory protein which is bound to the ligase in physiological context and present in the experimental structures. We refer to this experiment as ‘prediction with context’. Model choice was based on the ranking score provided by AF3.

### Model evaluation

To evaluate the similarity between the experimental structures and the predicted structures from AF-Multimer and AF3, the DockQ criteria were employed^30^. In short, DockQ provides a continuous score from 0 to 1 (with 1 being perfect similarity) which takes into account the fraction of conserved native contacts, RMSD of the target protein and the interface RMSD between a reference structure and a predicted structure^13^. We used the cutoff of 0.23 for acceptable models, in agreement with CAPRI criteria^31^. The choice of a relaxed cutoff is motivated by the fact that PROTAC systems are highly plastic and mobile^32^.

When predicting PROTAC-mediated interfaces with context (i. e., in presence of accessory proteins), only the DockQ score of the PROTAC-mediated interface was reported.

### Data availability

All the models analyzed in this article and the data used in Figure 2 are available in the Zenodo archive https://zenodo.org/records/14810843.

## Results

### Predicting Protein-Protein Interfaces using AlphaFold-Multimer

We collected 340 protein dimers not included in and not similar to the AF2 training set to evaluate the performance of AF2 on an unbiased set. Since PROTAC-mediated interfaces are both small and mediated by a ligand, we paid special attention to the interface size and the presence of ligand at the interface. Since very few interfaces are mediated by ligands in the test set, we also considered 20 ligand-mediated interfaces included in the AF2 training set. The predictions were evaluated by comparison with the experimental structures *via* the DockQ score, a continuous score in the range [0,1], that integrates the RMSD of the backbone of the shortest chain after superimposition of the longer chains, the RMSD of backbone interface atoms and the fraction of native interfacial contacts correctly predicted. A cutoff of 0.23 was used to discriminate correct from incorrect models, in agreement with CAPRI criteria^30,31^. In Figure 1, we show the model quality, assessed by the DockQ score, as a function of the size of the native interface, measured by ΔASA. Globally, AF2 produces correct models for 68% of the test cases that do not involve interface ligand (219 out of 316 back circles in Figure 1). In the presence of ligands (both open and filled red triangles in Figure 1), the proportion of correct models is equal to 85% (17 out of 20 open rend triangles and 4 out of 5 filled triangles in Figure 1). For the PROTAC-mediated complexes, the proportion of correct models drops to 18% (5 out of 28 green squares in Figure 1).

**Figure 1:**
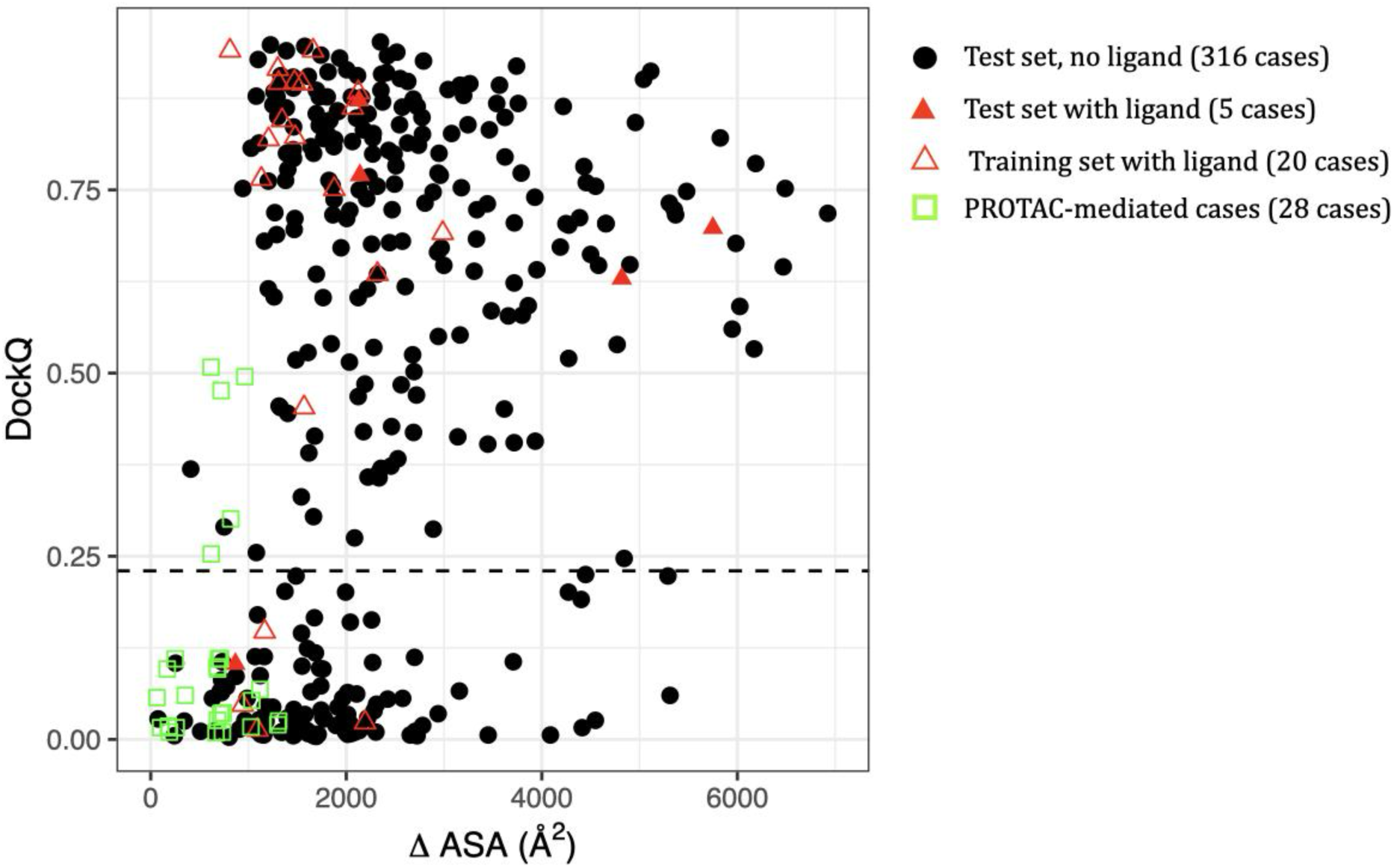
DockQ scores as a function of interface size. Here, the interface size is computed on the native complex. Black points: dimers from the test set with no ligand at the interface (316 cases). Filled Red triangles: dimers from the test set with ligand at the interface (5 cases). Open red triangles: dimers from the training set with ligand at the interface (20 cases). Open green squares: PROTAC-mediated dimers (28 cases). The horizontal dashed line indicates the threshold for acceptable models (DockQ=0.23).

As can be seen in Figure 1, the quality of the models predicted by AF2 is severely impacted by the interface size. Interfaces smaller than 1000 Å^2^ are very challenging to predict: the majority of the models have a DockQ score lower than 0.23 (average DockQ score<0.16, see Table S2) . There is a clear shift between 1000 and 2000 Å^2^, where about 60% of the models are of acceptable quality (average DockQ score 0.46, see Table S2), whereas for interfaces larger than 2000 A^2^, more than 75% of the models are of acceptable quality (average DockQ score >0.55, see Table S2). We observed a high correlation between the DockQ scores and the AF2 confidence scores, see Figure S1A, with a Pearson correlation coefficient equal to 0.79. However, some models of low quality have good AF2 confidence scores, see Figure S1B, particularly for small interfaces.

If we now focus on interfaces mediated by ligands (both open and filled red triangles in Figure 1), they seem to follow the same trend as other interfaces, with a strong influence of the interface size. PROTAC-mediated interfaces (green squares in Figure 1) are all small in size, and the predicted models are of poor quality. Except for five cases, the DockQ score is lower than 0.23. Increasing the sampling by AF2 with three seeds instead of one did not improve the results (Figure S2). We also monitored the influence of the ligand size, however, no clear trend could be identified due to the limited sample size. (Figure S3).

To further investigate the bias related to the interface size, we monitored the size of the predicted interfaces with respect to the native ones, in the case of wrong models (DockQ<0.23). The results are shown in Figure 2. We observe a general tendency of AF2 to predict models with larger interfaces than the native complex, when the interfaces to predict are small. Of note, interfaces smaller than 1000 A^2^, which are frequently mispredicted (see Figure 1), systematically produce models with much larger interfaces.

**Figure 2:**
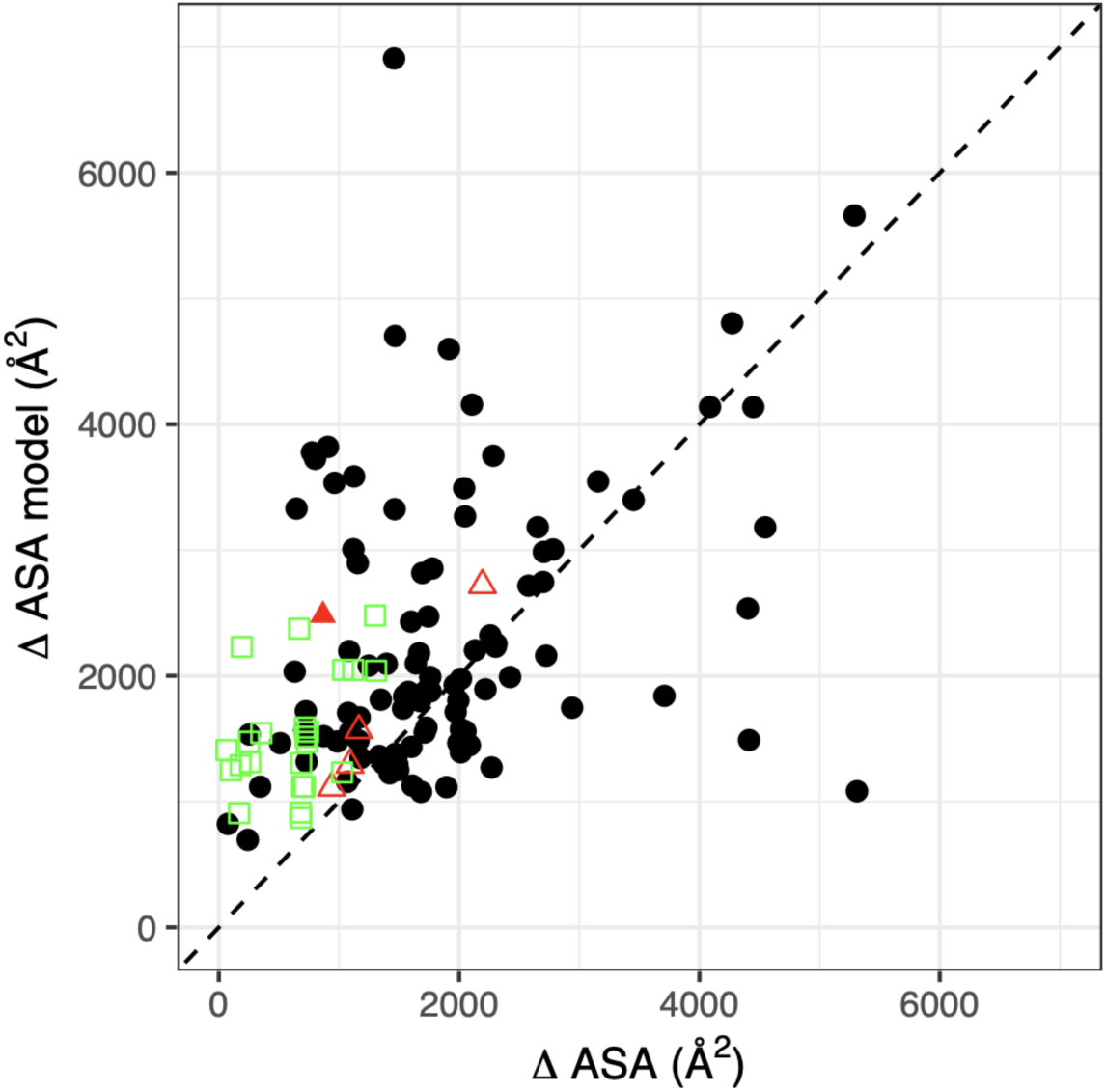
Interface size in the models as a function of the native interface size, for wrong models (DockQ < 0.23). Black points: dimers from the test set with no ligand at the interface (108 cases). Filled Red triangle: dimers from the test set with ligand at the interface (1 case). Open red triangles: dimers from the training set with ligand at the interface (4 cases). Open green squares: PROTAC-mediated dimers (23 cases).

In conclusion, the prediction of small interfaces with AF2 is still a challenge; and PROTAC-based interfaces typically belong to this category, preventing us from going further into the reasons of failure. On natural dimers, the presence of a ligand in itself does not seem to perturb the prediction.

### Attempted improvements of PROTAC-mediated interface prediction with AlphaFold 3

We tested the accuracy of AF2 and the newest AF3 on the set of challenging PROTAC-mediated dimers. The predictions were evaluated by comparison with the experimental structures *via* the DockQ score as explained before. The results are shown in Figure 3.

**Figure 3.**
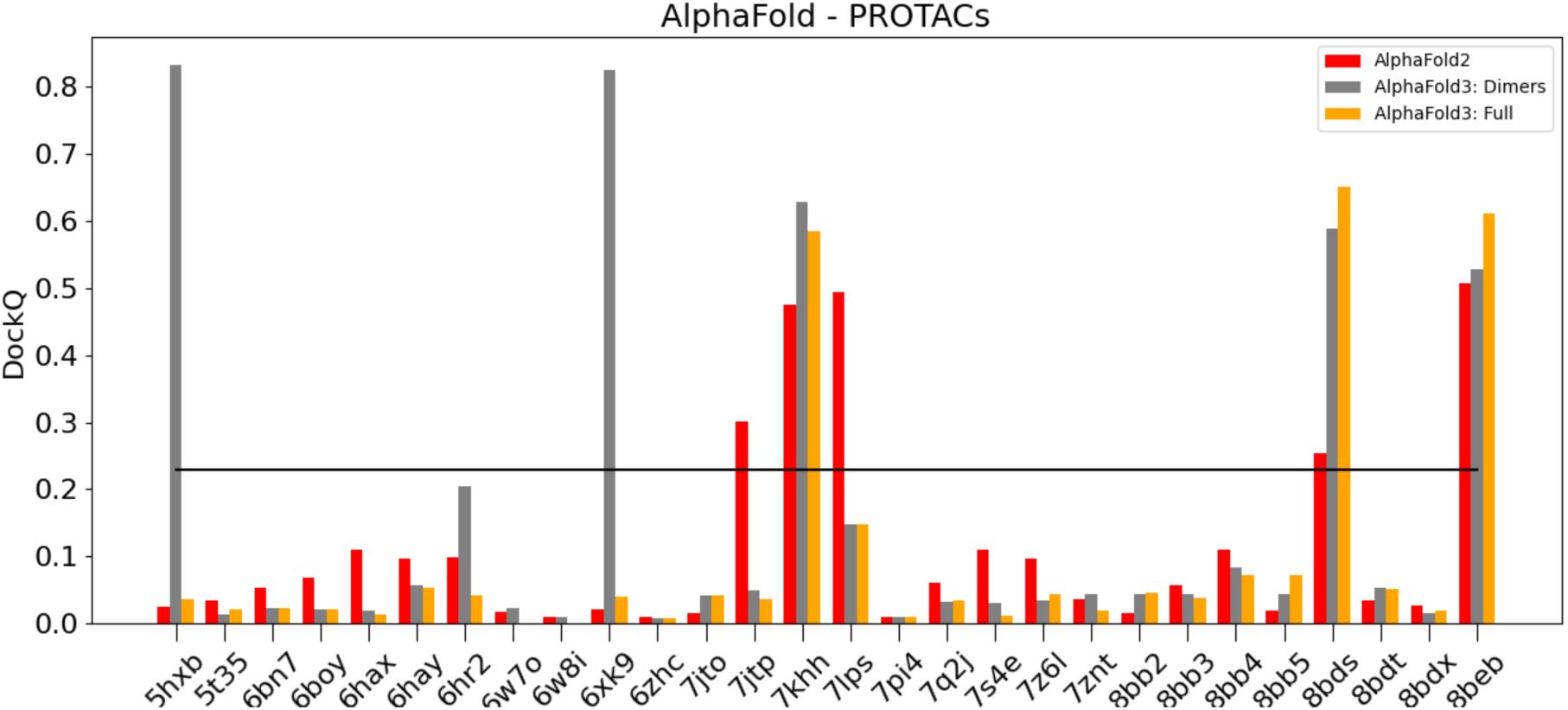
: Comparison of calculated DockQ values for all PROTAC-mediated systems using either AF2 or AF3 with respect to experimental crystal structures. The black line highlights DockQ’s acceptable criteria threshold at 0,23. For systems 6w7o and 6w8i, no accessory protein was present in the experimental structure.

As can be seen in Figure 3, AF3 generates acceptable models for 5 cases out of 28 (grey point in Figure 2). This is similar to AF2 global accuracy, which also generates 5 acceptable models (red points), although AF2 and AF3 do not succeed on the same cases (3 cases are correctly predicted by both). Of note, correct models generated by AF3 have relatively high DockQ scores (>0.5) compared to AF2, reflecting more accurate predictions.

We tested further the capability of AF3, by providing context to the prediction. Indeed, in physiological context the E3 ligase receptor is usually bound to accessory proteins, making a part of the surface inaccessible to an interaction with the target protein. We hypothesized that the addition of context, in the form of the accessory proteins found in experimentally determined structures, would improve the prediction of the receptor/substrate interface by limiting the search space available.

The E3 ligase receptor/accessory protein interfaces are highly conserved and should be easy to predict. However, the comparison between DockQ scores with and without context (orange *versus* grey bars in Figure 3) shows that the addition of context doesn’t improve the prediction: AF3 with context (AlphaFold3: Full, orange bars) generates acceptable models for 3 cases only *versus* 5 without context. Of note, these 3 cases were already well predicted by AF3 without context (Alphafold: Dimers, grey bars). Within these 3 cases with correct prediction with context, 2 cases have an improved DockQ score, although the increase is small. For example, the model for 8BDS has a DockQ score equal to 0.652 with context and 0.588 without context. For the large majority of systems, we find that the inclusion of the context does not significantly improve the prediction accuracy, and in some cases AF3 predictions including accessory proteins appear to be worse, as shown for 5HXB and 7Q2J in Figure 4. The predictions for all the 28 PROTAC cases are shown in Figure S5.

**Figure 4:**
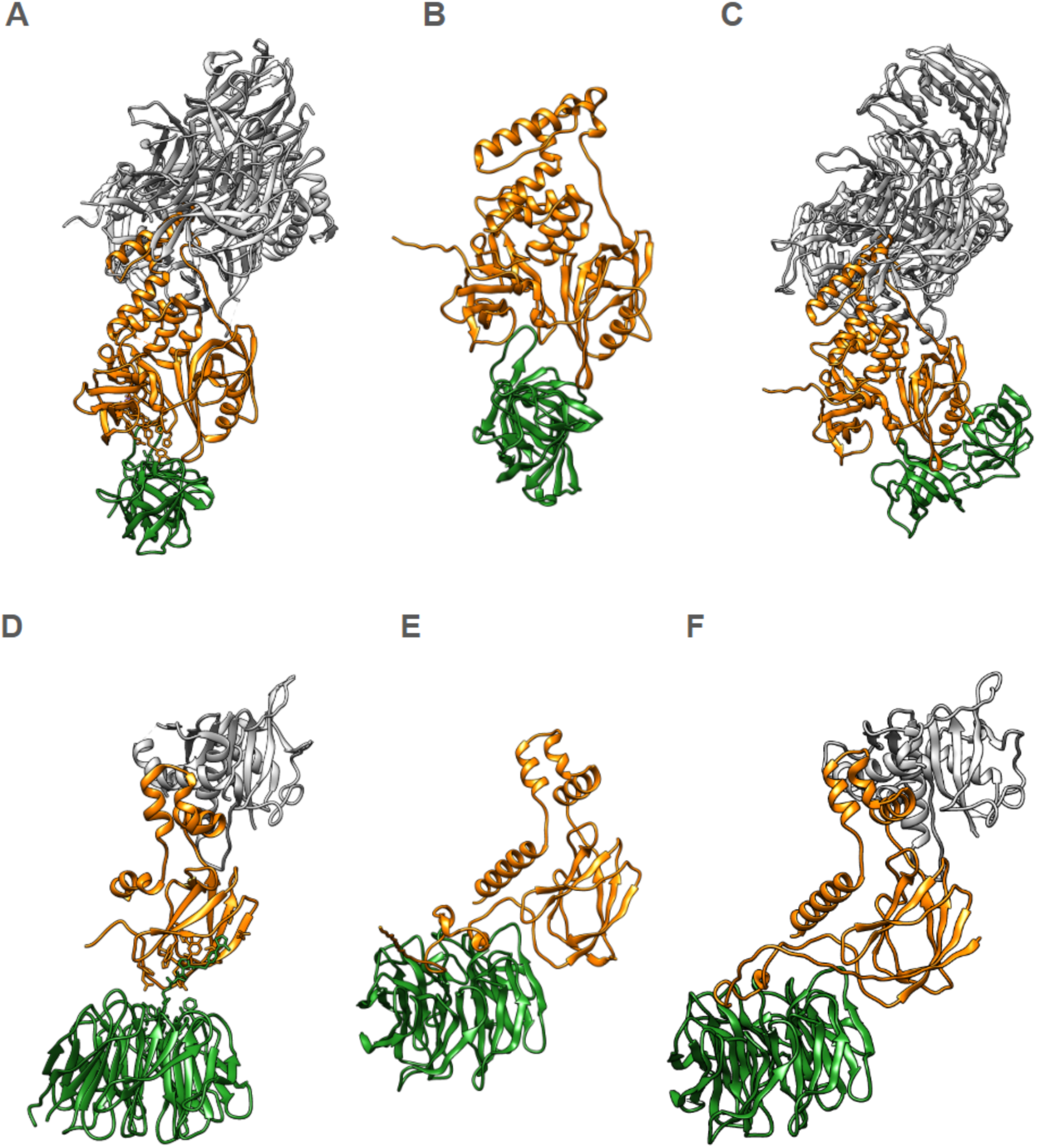
Top ranked predicted protein-protein interfaces for 5HXB (top) and 7Q2J (bottom). A,D) Crystal structures; B,E) AF3 models of dimers (DockQ: 0.833 and 0.032); C,F) AF3 models with inclusion of the accessory protein (DockQ: 0.036 and 0.043).

The inspection of models generated with context reveals that the interfaces between E3 ligase receptors and accessory proteins are generally well predicted, with all DockQ scores greater than 0.6 and a median DockQ score equal to 0.941, see Figure S4. However, this did not allow to improve the results for the E3 Ligase-Target interface. Given the current limitations of the AF3 web server in terms of available ligands, it was not possible to test whether inclusion of PROTACs would increase prediction accuracy. In conclusion, PROTAC-mediated interfaces remain very challenging to predict, even with AF3 and some context.

## Discussion

The prediction of E3-ligase/protein target complex can be the first step in computational pipelines for PROTAC design. In this context, the available information are the structures of E3 ligase in complex with a ligand, and of the protein target with a ligand. An important part of the PROTAC design is the optimization of the linker between the two ligands, which is determinant for the molecule’s catalytic activity^3,33,34^. In our previous work^13^, we developed a protocol to predict those complexes using LightDock^14^, a macromolecular docking framework that allows the incorporation of residue-based restraints. At that time, we also benchmarked AF2, with limited success, on PROTAC-mediated ternary complexes. A recent benchmark on a larger set of 43 PROTAC cases conducted by others and published during the preparation of this article confirmed the limitation of AF2 on PROTAC-mediated complexes^23^.

In the present study, we further investigate the reasons for this limitation. PROTAC-mediated interfaces have several peculiarities that could impact the prediction: (1) small interface size/area, (2) presence of ligand at the interface, (3) included in larger complexes with other proteins. Here, we tried to deconvolve these different factors. First, we show that AF2 suffers from a significant bias against small interfaces. Within the PPIs we studied, those with larger areas were typically better predicted than small ones (Figure 1). This bias was already present in the former version of AF2 trained on monomers, as observed by Yin et al^35^ who benchmarked AF v.2.0 with the residue index hack on the 152 complexes of the Protein-Protein Docking Benchmark 5.5 ^36^. Our results indicate that this bias persists in the version of AF2-multimer that was specifically trained on protein-protein complexes. More sophisticated approaches that build upon AF like AFSample^37^ (that uses dropout during inference) and AFProfile^38^ (that denoises the input MSA) can improve the prediction in some cases. The potentiality of these approaches in the case of small interfaces remains to be explored. Another alternative would be to fine-tune AF for small interfaces.

Second, concerning the presence of a ligand at the interface, we have considered protein-protein interfaces that involve ligands for more than 10% of their size, to see if we could predict them in the absence of ligands - since AF2 does not model ligands. Our results indicate that these interfaces do not pose a specific challenge for the prediction, as many of them were predicted with DockQ scores between moderate (0.49) and high quality (0.80). However, both ligand-mediated and non-ligand-mediated interfaces appear to be ubiquitously affected by the interface size problem. Because all the known PROTAC-mediated interfaces are small, we cannot disentangle the effect of interface size from an hypothetical PROTAC-specific effect, as it would require a comparison with large PROTAC-mediated interfaces.

Third, despite the limitation of AF2 on PROTAC-mediated complexes, we also tested the effect of providing some context to the prediction: by adding the accessory proteins, the interface accessible to the interface with the substrate would be more limited, narrowing down the solution space. This was done with AF3. However, we noted no improvement in the predictions, with or without context.

A known peculiarity of PROTAC-mediated complexes is that they are induced and stabilized by the ligand, and the two proteins would probably not interact strongly in the absence of the PROTAC molecule. The resulting absence of an evolutionary signal could be another source of challenges for the accurate prediction of these complexes. A recent work of Roney and Ovchinnikov^39^ provides evidence that AF2 indeed learned an implicit energy function that encapsulates the physics governing the folded state and that the evolutionary signal serves as a guide in the global search for the optimal structure. Another recent study leads to the same conclusion, observing that AF has a remarkable capability to recover correct structures from certain perturbations without additional information provided by the multiple sequence alignment.^40^ Along the same line, AF2Complex, proposed by Gao et al^19^, successfully predicted hetero-dimers starting from unpaired multiple sequence alignments, suggesting again that the AF2 network learned an implicit energy function that is enough to represent the physics of protein-protein interfaces. So, in theory, it should be possible for AF2, and by extension AF3, to predict a non-natural interface in the absence of an evolutionary signal.

The absence of the PROTAC molecule likely poses a significant challenge, as the PROTAC molecule itself has a key role in stabilizing the protein-protein interface of these complexes. Given that these are large and flexible ligands, any algorithm that explicitly models the ligand must be capable of sampling (or receiving as input) multiple ligand configurations and then generate an ensemble of ternary complex configurations. Ensemble-generation methods could then be used to maximize the likelihood of finding physiologically relevant structures^32,41–44^.

The release of RoseTTAFold-All-Atom^32,45^ in 2024, might offer an alternative to the AF family of methods, as it is able to consider the effects of arbitrary ligands within protein-protein interfaces. It remains to be seen how well this modeling suite achieves in predicting highly plastic, “artificial” and shallow protein-protein interfaces such as those in PROTAC-mediated systems. Methods like AlphaLink, could get around the problem of ligand modeling by the incorporation of restraints in the prediction network^46^.

To summarize, because of their small size and peculiar physics, PROTAC-mediated complex prediction remains challenging even with cutting-edge deep-learning methods. Potential solutions to overcome these limitations include the development of models trained on small interfaces, and models that better consider the effects of ligands or incorporate restraints. Additionally, training models on conformational ensembles extracted from molecular dynamic simulations of PROTAC-mediated complexes is likely to yield better predictions for PROTAC-mediated PPIs. Since the field of deep-learning based structure prediction is rapidly evolving, new ligand-aware models have been developed during the course of the present study. Most recent methods like Chai-1^47^ and Boltz-1 remain to be tested; they could change the way PROTAC design is traditionally addressed since they allow the direct inclusion of ligands in the prediction^46,48^. To put our result in a broader perspective, the limitation of AF for the prediction of complexes with small interfaces calls for caution when working with multimeric systems that likely involve small interfaces.

## Supporting Information

Table S1: PDB Ids used in the study; Table S2: Repartition of cases by interface size range and model quality; Figure S1: Model quality, AF2 confidence score and interface size plots for the full dataset; Figure S2 : Model quality and AF2 confidence score in the PROTAC set, with 1 and 3 seeds; Figure S3. DockQ score as a function of ligand molecular weight; Figure S4 : AlphaFold3 results for the 28 PROTAC-mediated complexes; Figure S5 - AlphaFold Models for the 28 PROTAC cases.

## AUTHOR CONTRIBUTIONS

The manuscript was written and revised by all authors. Approval for publication was obtained from all authors.

## ACKNOWLEDGMENTS

G.P, C.G, P.C.T.S and J.M would like to thank the support of the French National Center for Scientific Research (CNRS). G.P. and P.C.T.S would also like to thank the funding from research collaboration agreements with PharmCADD. We also acknowledge the support of the Centre Blaise Pascal’s IT test platform at ENS de Lyon (Lyon, France) for the computer facilities. The platform operates the SIDUS solution developed by Emmanuel Quemener^49^. For the purpose of open access, the authors have applied a Creative Commons Attribution (CC BY) license to any Author Accepted Manuscript version arising from this submission.

## ABBREVIATIONS

AF: AlphaFold
PPI: Protein-Protein Interfaces
PROTAC: Proteolysis Targeting Chimera
MSA: Multiple Sequence Alignment

## Supplementary Information

**Table S1.**
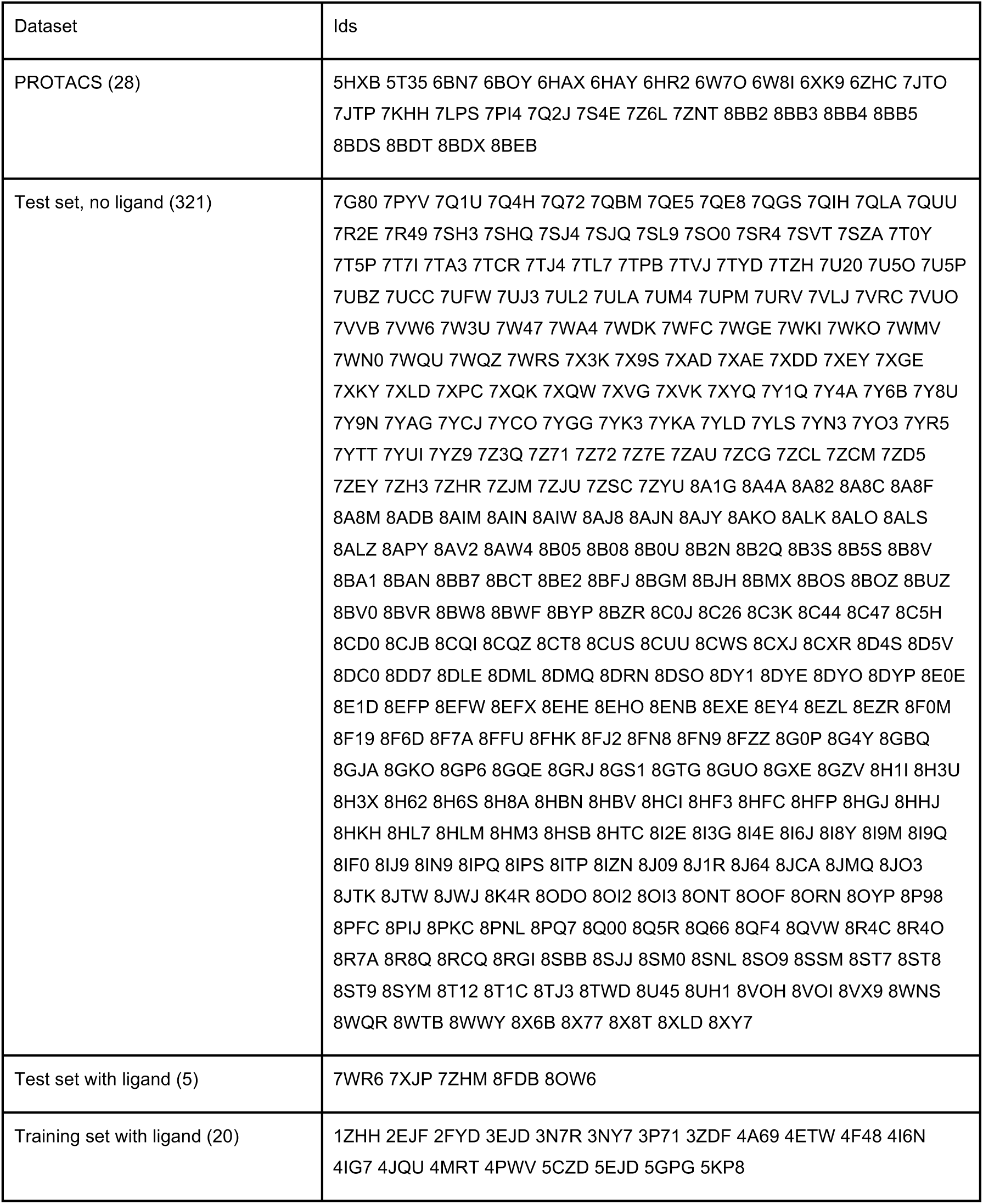
PDB Ids used in this study. Numbers in parentheses refer to the number of cases.

**Table S2.**
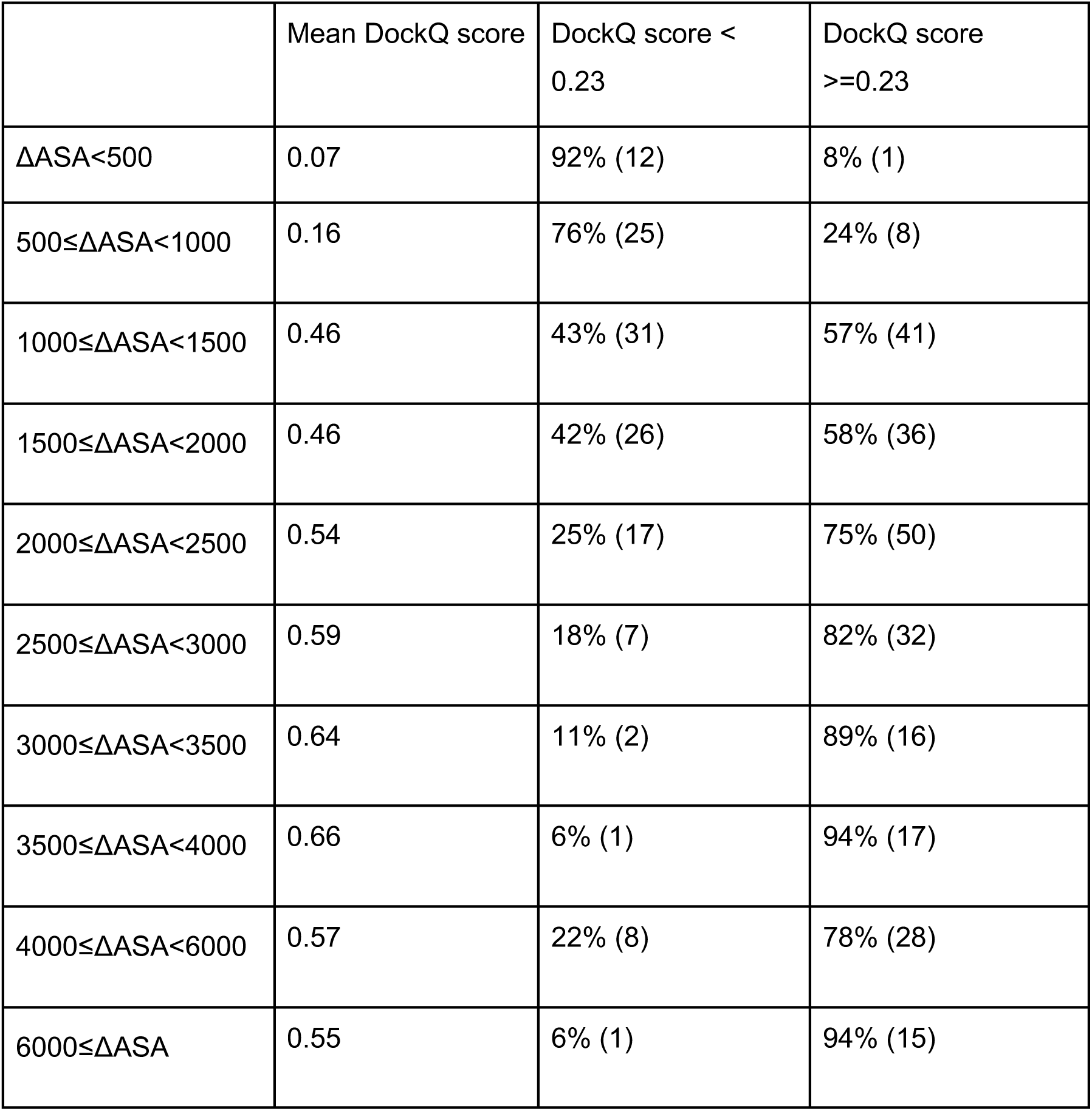
Repartition of cases by interface size range (expressed in Å^2^) and model quality (assessed by the DockQ score). Numbers in parentheses correspond to numbers of cases.

**Figure S1.**
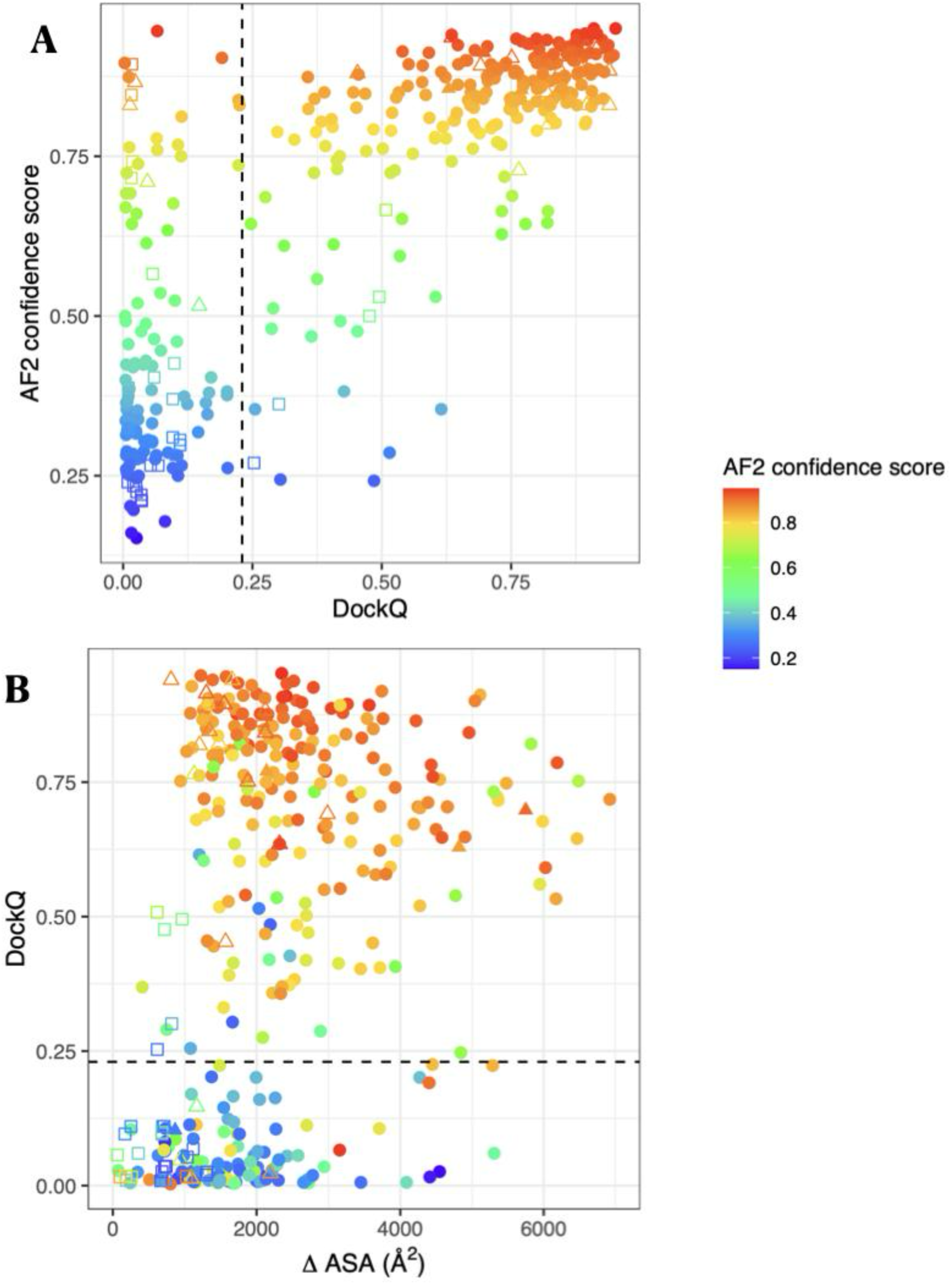
Model quality, AF2 confidence score and interface size. Points are colored according to the AF2 confidence score (0.8 ipTM+0.2pTM).

**Figure S2.**
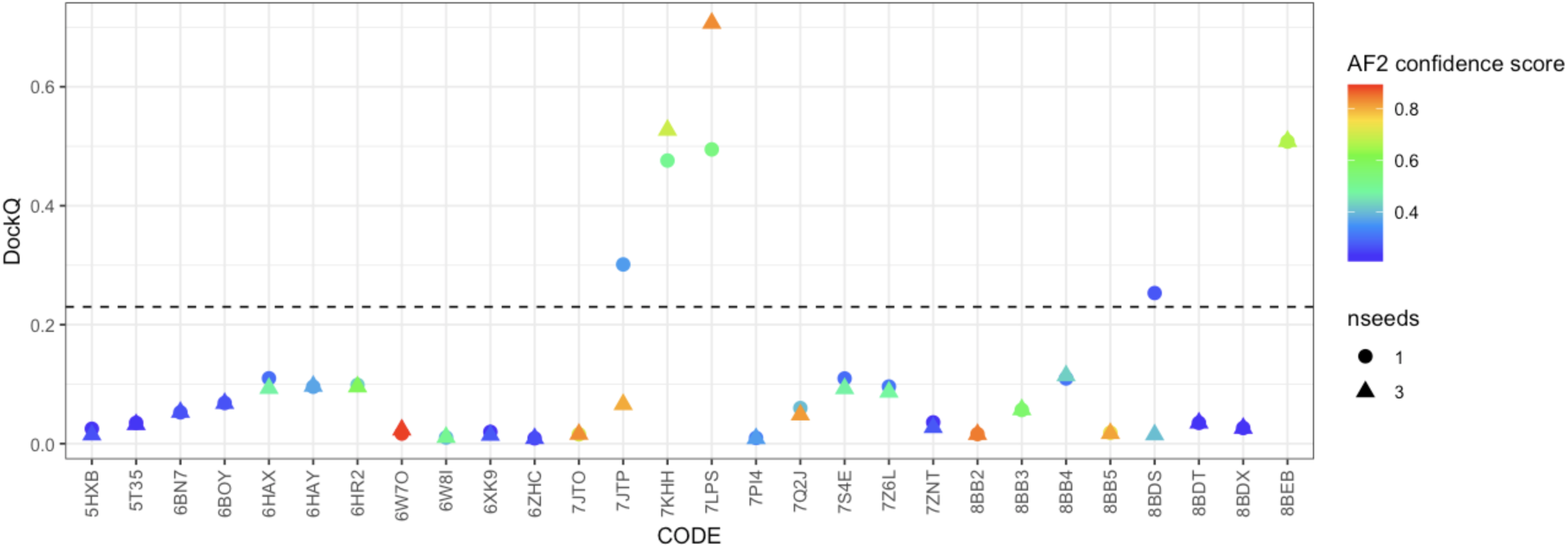
Model quality and AF2 confidence score in the PROTAC set, with 1 and 3 seeds. Points are colored according to the AF2 confidence score (0.8 ipTM+0.2pTM), and the shape indicates the number of seeds used for AF2 calculation.

**Figure S3.**
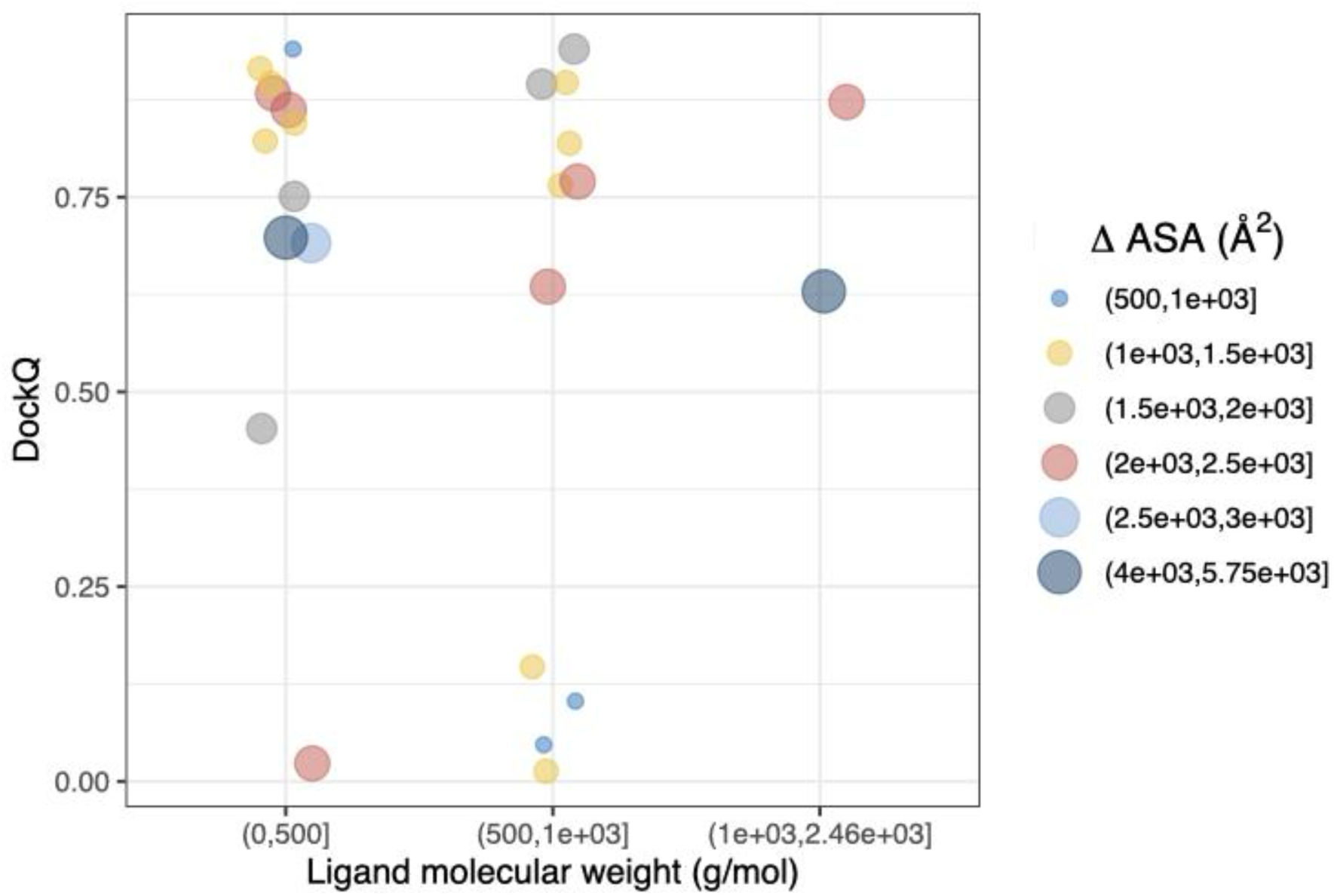
DockQ score as a function of ligand molecular weight for the 25 ligand-mediated complexes (not PROTACs-mediated complexes). Four out of five wrong predictions are for interfaces mediated by ligands with high molecular weight (>500g/mol). Note that these wrong predictions also involve small interfaces (small circles), and that the majority of complexes involving ligands of high molecular weight are well predicted.

**Figure S4.**
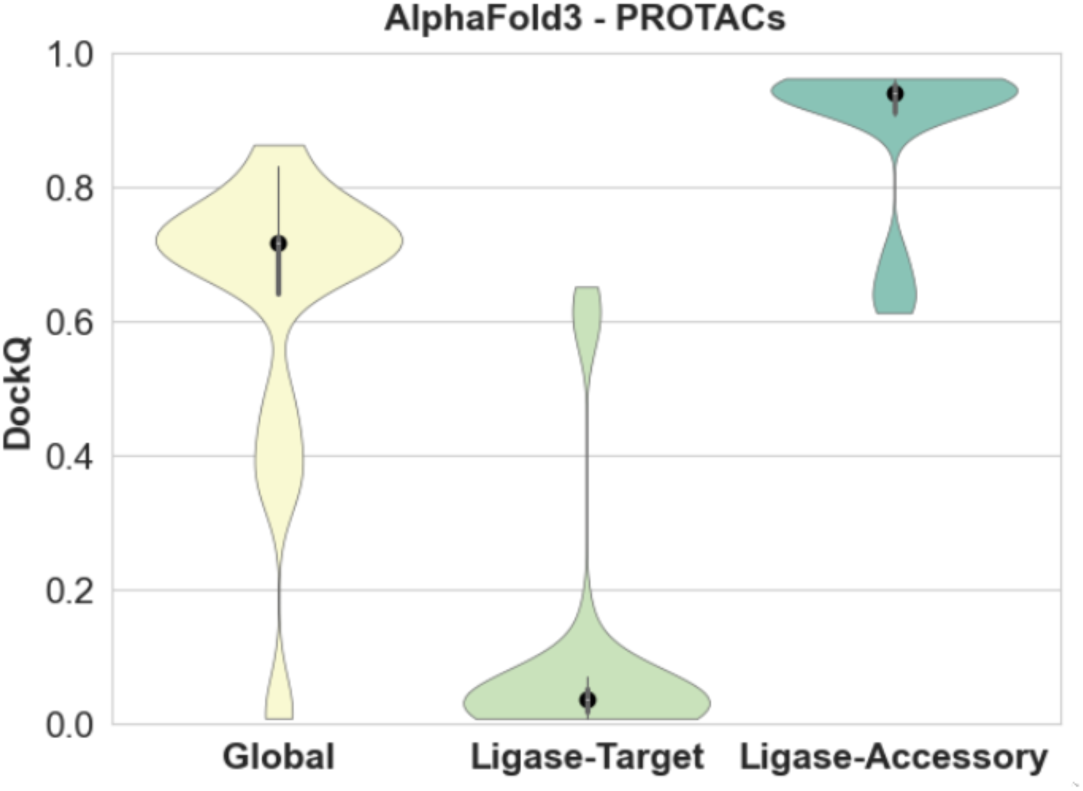
AlphaFold3 results for the 28 PROTAC-mediated complexes. In this experiment, all proteins within the PDB entry were provided to AlphaFold server and all 5 top-predicted models were queried against their corresponding crystal structure. We extracted the best ranked solution to AF3.

The data in Figure S4 show that AF3 is apparently performing at an acceptable level for the full systems, with an average DockQ score of 0.75. However, once these predictions are separated into two categories, the Ligase-Target interface DockQ and the PPIs involving the ligase and accessory proteins, we see the same pattern as in AF2-Multimer. In general, Ligase-Target interfaces remain difficult to predict with AF3, with only 4 systems out of 28 achieving a DockQ score above 0.23 for at least one predicted structure. This result is only marginally better than the result obtained with AF2, where three out of 28 systems were correctly predicted. In contrast, all interfaces not including the target protein were well predicted (all above Dock > 0.6) and the average DockQ score was around 0.9. This data paint a concerning picture: While on the surface these multi-protein complexes appear to be correctly predicted by AF3, it turns out that the most important interface is still missed in the large majority of the cases.

**Figure S5.**
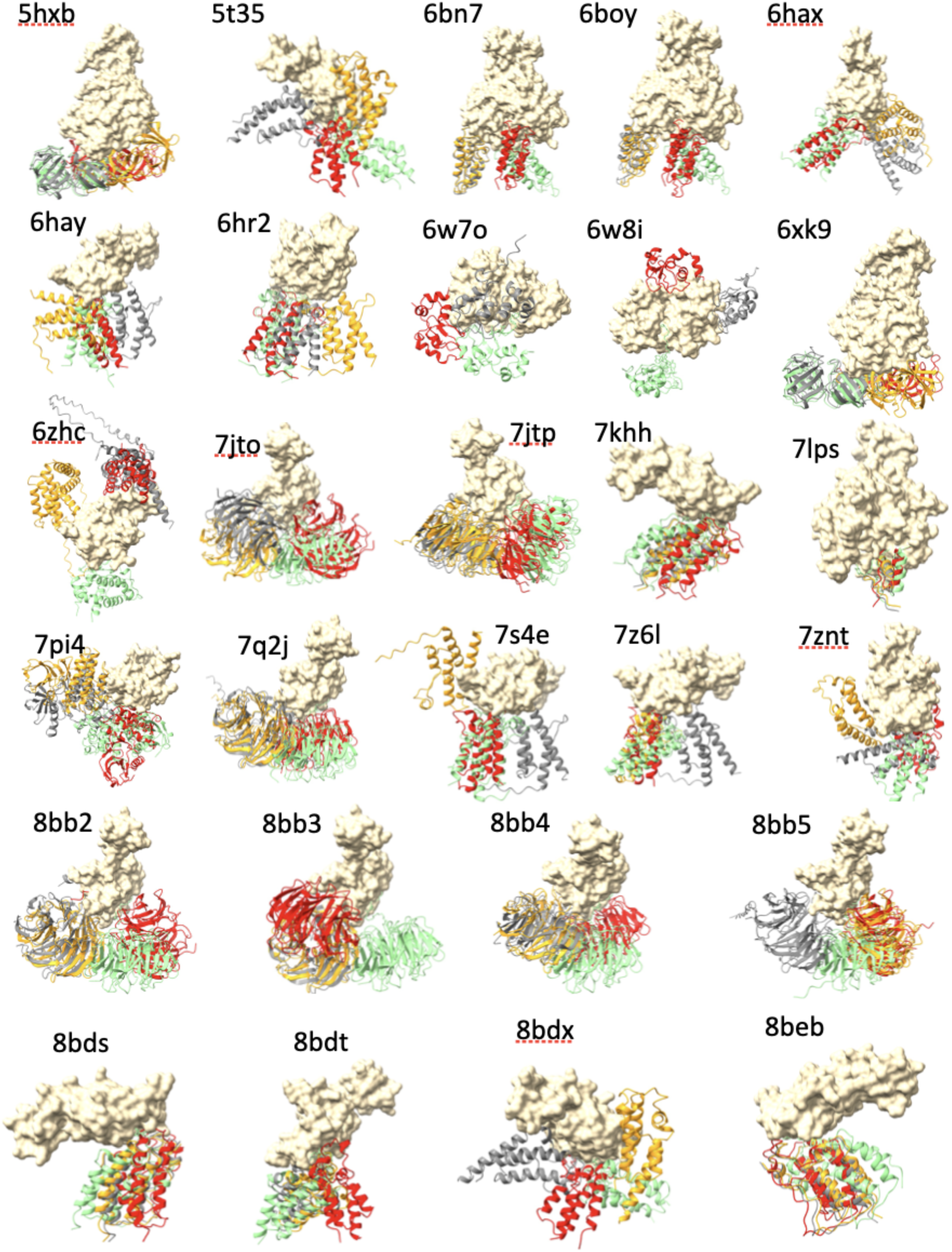
AlphaFold Models for the 28 PROTAC cases. The E3 ligases are shown in surface representation and the protein targets in cartoon representation with the following color code: green=experimental structure, red=AF2 prediction, gray=AF3 prediction, orange=AF3 prediction with context.

